# Detecting deception in event narratives: the analysis of response latencies and error rates to unexpected questions

**DOI:** 10.64898/2026.10.29.762734

**Authors:** Maria Vittoria Rosa, Giulia Melis, Giuseppe Sartori, Andrea Zangrossi

**Author notes:** Corresponding author (MVR).

## Abstract

Deception is a central phenomenon in the study of human behaviour, and in forensic and investigative contexts it assumes significance due to its potential consequences. Although the unexpected questions technique has shown promising results, its application to the narration of a false autobiographical event remains underexplored. This study aims to fill this gap using the unexpected questions technique in an investigative interview setting. A total of 60 participants were randomly divided into two groups: the honest group was asked to describe their most recent day trip, while the deceptive group was asked to memorise a narrative describing a day trip. Both groups underwent an audio-recorded interview containing control, expected and unexpected questions. The results showed that both groups exhibited longer response latencies and higher error and inaccuracy rates in response to unexpected questions, but these indices were significantly higher in the deceptive group (*p = .022* for response latency, uncorrected for multiple comparisons; *p < .001* for error and inaccuracy rates). The findings suggest that longer response times and higher error and inaccuracy rates during unexpected questions may reflect not only the difficulty of the questions, but also additional cognitive demands faced by deceptive participants. Furthermore, these indicators were used to classify a subject as deceptive or honest using machine-learning algorithms. Within the sample, the model showed excellent cross-validated discriminatory performance (AUC = 0.967), suggesting that these indices may have diagnostic potential worthy of further investigation.

## Introduction

It has been estimated that people lie on average once or twice a day during social interactions [1], yet the human ability to reliably detect deception is very limited, with an accuracy of approximately 50% in distinguishing between truth and deception [2]. While casual lies (low-stakes lies) permeate daily interactions, the focus of the present research specifically targets the detection of prepared false event narratives within an interviewing context, which involve greater risks (high-stakes lies) [3–6]. The discrepancy between deception frequency and the insufficient human ability in detecting lies creates a profound practical problem in forensic settings, where testimony traditionally represents one of the main sources of evidence. Parallel to this practical issue is a crucial theoretical problem: despite the variety of lie-detection techniques available, many have limitations in terms of applicability or reliability [7–13]. Although indirect cognitive indicators such as response latency show promising utility, their diagnostic efficacy is continuously undermined by the effects of deception prepared in advance.

This study is based on the Activation-Decision-Construction-Action Model, which posits that deception generally requires more controlled processing than truth telling, which more often relies on direct retrieval from memory [14–16]. This increased cognitive load manifests itself in several indicators, among which response latency has emerged as one of the most promising in deception research [16–20]. However, deception prepared in advance, such as an alibi, is already stored in long-term memory, resulting in an alteration of cognitive demands which leads to shorter response times compared to spontaneous lies [15, 16, 21, 22].

To address this limitation, Vrij and colleagues [23] developed the unexpected questions technique, which is designed to reduce the benefits of advance preparation. In response to these questions, honest subjects must retrieve information that may not have been recently rehearsed, whilst deceptive participants are additionally forced to construct new details consistent with their false narrative. To distinguish between honest and deceptive subjects, response latencies to unexpected questions were compared with the baseline, consisting of response latencies to expected and control questions. The effectiveness of this technique has been supported by several studies [19, 24–31]. More recently, a semi-systematic review of the field concluded that the unexpected questions paradigm represents one of the most extensively investigated cognitive approaches to deception detection, although the evidence remains heterogeneous and further replication studies are needed [32]. Research conducted by Monaro and colleagues in 2017 [33] showed that analyses based on expected questions correctly distinguished deceptive from honest participants with accuracies ranging from 65% to 67%, whilst classification based on unexpected questions achieved an accuracy of 95%. Furthermore, a meta-analysis of cognitive lie-detection approaches (imposing cognitive load, encouraging interviewees to say more, and asking unexpected questions) reported 71% overall accuracy, compared with 56% for standard approaches [34]. More specifically, Melis et al. [18] reported 98% accuracy (mean false positives rate= 0.1; means false negative rate= 0) in classifying participants as honest or deceptive using a machine-learning model based on response latencies and error rates across the three question types.

Despite the particular importance of event narratives in forensic contexts, most research projects have focused on identity deception rather than the deception of autobiographical events [18, 19, 33]. Recounting a false and complex autobiographical event requires access to episodic memory and the integration of coherent details, which is likely to demand more cognitive resources than reciting a fabricated identity. In the present study, this demand was operationalised by contrasting genuine episodic retrieval (honest group) with the retrieval of a recently memorised false event narrative (deceptive group). The deceptive task therefore primarily taxes short-term memory for a prepared script and the on-the-spot inference of details that the script does not state, rather than the continuous monitoring of a self-generated false episodic memory. Furthermore, most lie detection studies are based on the bald-faced lie (BFL) vs. bald-faced truth (BFT) dichotomy, requiring the participant to tell either a 100% lie or a 100% truth [22, 35, 36]. Instead, real investigative interviews involve a wide variety of lies and the distortion or omission of the truth rather than a completely crafted lie, which would require more cognitive resources [37, 38]. Critically, most previous studies have often designed their research framework around features which do not adequately reflect investigative settings, limiting their applicability to real-world contexts [22].

This study follows the foundational steps laid out by Melis et al. [18], measuring response latencies and error rates in response to unexpected questions in a setting similar to an investigative interview. Our intent is to further investigate the performance of the unexpected questions technique in the recognition of deception of an autobiographical event. Furthermore, we strive to overcome the bald-faced lie vs. bald-faced truth dichotomy and to move toward greater ecological validity by introducing the metric of “inaccuracy rate”. This addition was intended to capture not only wrong or slow responses, but also whether participants adopt less precise response strategies to avoid errors. Note that participants were still assigned to recount either a wholly true or a wholly fabricated (memorised) event; the departure from the dichotomy therefore concerns the coding of responses (correct, erroneous, inaccurate or omitted) rather than the generation of the narrative.

## Methods

### Design and hypothesis

Based on what has been said so far, the manipulated independent variables included the question type (unexpected vs expected vs control) and the participant’s veracity (honest vs deceptive). Deception was tested under a specific task configuration, as described below. The dependent variables measured were response latency, error rate and inaccuracy rate. To this end, we hypothesize that for both groups answering unexpected questions will result in longer response latencies and higher error and inaccuracy rates, due to the additional difficulty of adapting premeditated responses on the spot. Additionally, we predict that deceptive participants will show longer response latencies and higher error and inaccuracy rates when answering expected and unexpected questions compared to honest participants, due to the greater cognitive load required. Finally, we expect that answering unexpected questions will be perceived as more cognitively demanding and less predictable than other question types.

### Participants

The study examined a sample of 60 Italian participants aged between 19 and 29 (M= 23.97, SD=2.09), of whom 32 were female (53.3%) and 28 were male (46.7%). The only inclusion criterion was being a native Italian speaker. It has previously been demonstrated that a sample of 60 participants is sufficient to detect a medium-sized effect (effect size = 0.25), with a 95% probability ((1-β) = .95), using a two-group mixed ANOVA targeting the within–between interaction, whilst maintaining the error rate at 5% (α = .05) [18].

### Experimental procedure

Ethical approval was obtained from the Ethics Committee of the Department of General Psychology at the University of Padua. Written informed consent was obtained from all participants before the experiment began; moreover, they were advised of the confidentiality of their data and their right to withdraw at any time without repercussions. The research was conducted according to the principles expressed in the Declaration of Helsinki. All material provided to participants was in Italian and the interviews were conducted in Italian. The recruitment period and participant data collection took place from 10 October 2025 to 9 December 2025.

An online questionnaire was administered in person to collect demographic information (gender, age, educational level and occupation) and to assign two conditions to the participants using a computer-generated randomisation sequence, with block randomisation (block size = 2). Participants were asked either to memorise a short story about a recent day trip, classifying it as belonging to the deceptive group (n = 31), or to describe their own most recent day trip, placing it within the honest group (n = 29). Two participants were excluded from the analyses because they had misunderstood the task instructions, and two replacement participants were subsequently recruited and assigned to the experimental conditions using Qualtrics. The final sample consisted of 60 participants: 31 in the deceptive group and 29 in the honest group. Before writing their account, participants in the honest group were instructed to focus on specific elements of the event to establish a clear reference narrative for subsequent response coding. The deceptive group was given five minutes to familiarise themselves with the provided story, before performing a free recall task designed to ensure complete, error-free memorization. Provided that no errors had been made, the deceptive participant moved on to the question phase, otherwise, they were given another five minutes to memorise the story and retell it again. All participants were then audio-recorded and interviewed on the day trip story they had either memorised or described. The questions were categorised into 12 control (C) (*e.g. Which city are you in?*), 12 expected (E) (*e.g. What time did you wake up?*) and 12 unexpected items (U) (*e.g. How long did it take you to reach your destination?*), and were presented in randomized order. This random layout was used as a task-switching method implemented to simulate high-stakes lies by increasing the cognitive load. One aspect of the design that should be considered in interpreting the findings is that the honest group answered questions about a self-experienced event, while the deceptive group answered questions about a recently memorised story. Therefore, the study examined how deception manifests within a specific task framework. In the end, participants were asked to complete two online questionnaires, administered in person, regarding their perception of the difficulty in answering each question and the level of predictability of the questions. Both perceptions were measured on a Likert scale from 1 (being “*very easy*” / “*unpredictabl*e”) to 5 (being “*very hard*” / “*very predictable*”). The experiment lasted approximately 20 minutes. The entire process of conducting the interviews and analysing the audio-recordings was entrusted to a single experimenter; therefore, the double-blind procedure was not adopted. The list of questions and the materials used can be found in the supplementary materials (S15 File 1).

### Audio-analysis

Audacity Software (https://www.audacityteam.org/) was used to extract response latency, i.e. the response time (RT) measured from the moment the interviewer finishes the question to the moment the subject begins to provide the answer, considering the first meaningful word and excluding filler words such as “*umm*”. Each response was compared with the reference narrative and coded as correct, erroneous, inaccurate or omitted. Responses were categorised as *inaccurate* if they: a) contained additions not specified or inferable from the text; b) did not provide the precise answer but rather the calculations leading to the answer; c) did not provide the precise and specific data as requested, instead circling around the answer; d) contained only part of the correct answer, omitting some data. Conversely, responses were categorised as *omissions* if the subject replied a) that they did not fully recall the answer, i.e., when not even part of the answer was provided; b) “*I don’t know*”. Cases in which the subject provided an incorrect or garbled answer were classified as “*errors*”.

### Statistical analysis

Prior to analysis, the mean scores for response latency, perceived difficulty and perceived predictability were calculated across the three question conditions. The error and inaccuracy rates were calculated using binary coding (error rates: error= 1, every other response=0; inaccuracy rates: inaccuracy= 1, every other response= 0) and calculating the average scores for the three question conditions.

All statistical analyses were conducted using JASP (Version 0.95.4; https://jasp-stats.org/). A significance threshold of α = .05 was applied. Greenhouse-Geisser correction was used for sphericity violations. The Welch t-test was used for unequal variances between the groups; an asterisk in the table denotes the specific statistical test (Welch or Student) employed for each condition. Effect sizes were reported as partial η²*_p_* for ANOVAs and Cohen’s d for t-tests. Bonferroni correction was applied to post-hoc pairwise comparisons among the three question types following the repeated-measures ANOVAs. No multiple-comparison correction was applied to the between-group independent-samples t-tests.

### Machine-learning analysis

Two machine-learning models were constructed through PyCaret implemented in Python. Logistic regression and random forest models were developed, optimised by tuning the hyperparameters via grid search with 10-fold cross-validation, to identify the parameter configuration that maximised predictive performance and reduced the risk of overfitting. To construct these models, the group variable was transformed into a binary variable, where the positive class (1) corresponds to deceptive and the negative class (0) to honest individuals. The response features used for classification were latency, error rate and inaccuracy rate. For the subject-level classification, the three indicators were computed for each of the three question conditions (control, expected and unexpected); for the item-level classification, they were computed for each individual response to an unexpected question (720 responses: 60 participants × 12 unexpected questions). For each machine-learning algorithm, various performance metrics were calculated: accuracy, area under the curve (AUC, i.e. how well the model separates the two classes), precision, recall and F1 score.

## Results

The following results show the effects of veracity (honest vs deceptive) and question type (control vs expected vs unexpected) on response latency, error and inaccuracy rates, perceived difficulty and predictability, and classifications at a subject and item level.

### Analysis of response latency

To examine the impact of veracity and question type on response latency, a 2 (veracity: honest vs. deceptive) x 3 (question type: control vs. expected vs. unexpected) mixed ANOVA was conducted. The analysis, in which the Greenhouse-Geisser correction was used, demonstrated significant effects for question type, F*_GG_*(1.277, 74.080) = 107.783, p < .001, *η^2^_p_* = .650, and for the interaction between veracity and question type, F*_GG_*(1.277, 74.080) = 8.355, p = .003, *η^2^_p_* = .126. The main effect of veracity was not significant, F*_GG_*(1.277, 74.080) = 1.601, p = .211, *η^2^_p_* = .027. These results indicate significant associations of question type, and of the interaction between this factor and veracity with response latency. Furthermore, an independent samples t-test was conducted to explore the effect of all three question types on response latency. The results (Table 1) revealed a significant difference between the two groups in response latencies for unexpected questions, *t_student_*(58) = −2.347, p = .022, d = −0.606, whilst no significant differences emerged for control questions (p*_student_* = .655) or expected questions (p*_welch_* = .078). These results partially support our initial hypothesis, by showing that unexpected questions increase response latency in both groups, with a significantly greater effect in the deceptive group.

**Table 1.**
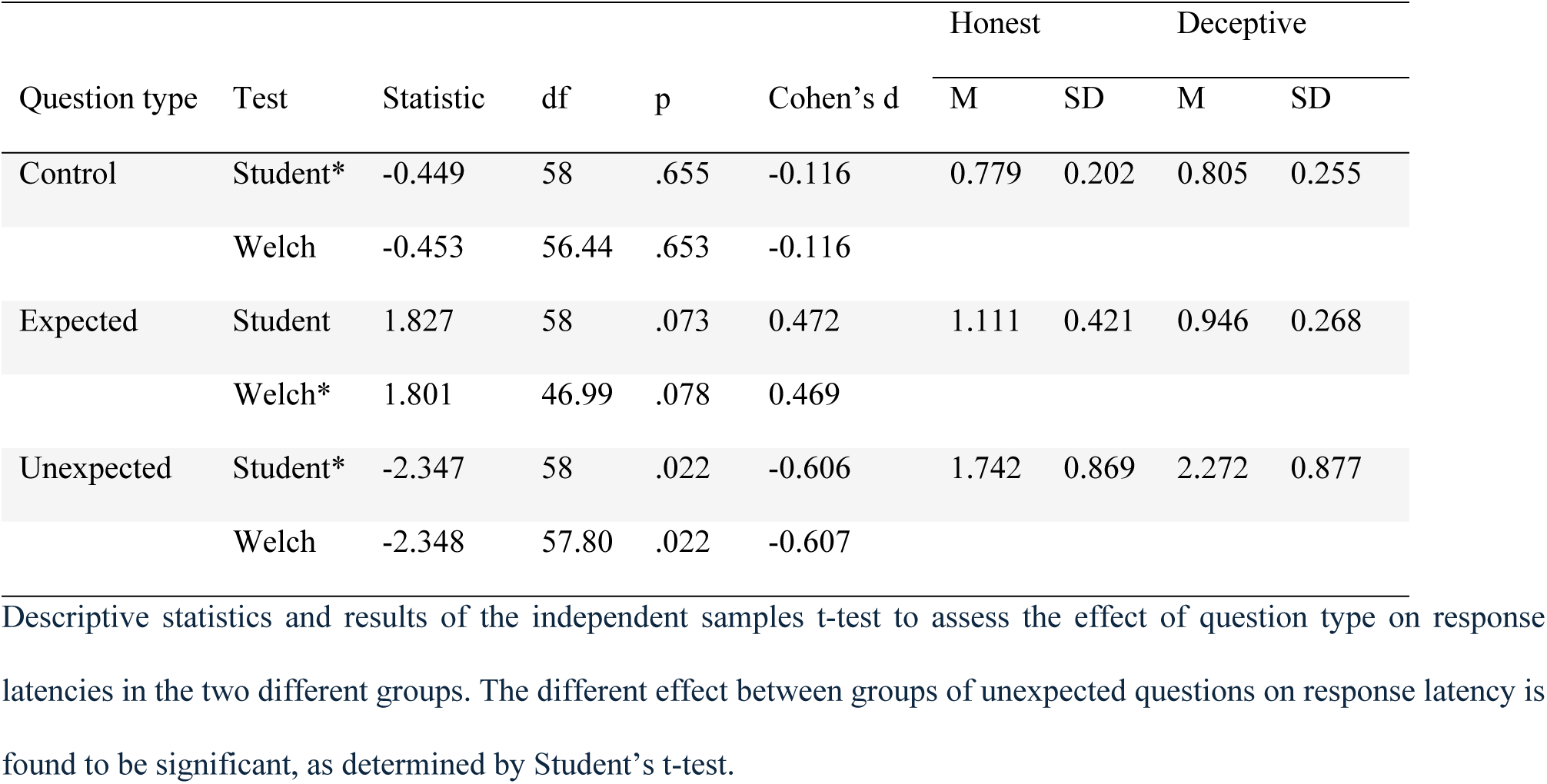
Independent samples t-tests and descriptive statistics of response latencies.

|  |  |  |  |  |  | Honest |  | Deceptive |  |
|---|---|---|---|---|---|---|---|---|---|
| Question type | Test | Statistic | df | p | Cohen’s d | M | SD | M | SD |
| Control | Student* | -0.449 | 58 | .655 | -0.116 | 0.779 | 0.202 | 0.805 | 0.255 |
|  | Welch | -0.453 | 56.44 | .653 | -0.116 |  |  |  |  |
| Expected | Student | 1.827 | 58 | .073 | 0.472 | 1.111 | 0.421 | 0.946 | 0.268 |
|  | Welch* | 1.801 | 46.99 | .078 | 0.469 |  |  |  |  |
| Unexpected | Student* | -2.347 | 58 | .022 | -0.606 | 1.742 | 0.869 | 2.272 | 0.877 |
|  | Welch | -2.348 | 57.80 | .022 | -0.607 |  |  |  |  |

In addition, two separate repeated-measures ANOVAs were conducted to examine the effect of question type within each group. The Greenhouse-Geisser correction was used in the analyses. The results for the deceptive group revealed a significant effect of question type, F*_GG_*(1.227, 36.810) = 75.06, p < .001, *η^2^_p_* = .714. A post hoc analysis revealed significant differences between responses to control questions compared with unexpected ones (p*_bonferroni_* < .001) and between expected questions compared with unexpected ones (p*_bonferroni_* < .001). No differences were found between the control and expected conditions (p*_bonferroni_*= .086). The same pattern was observed in the honest group, F*_GG_*(1.216, 34.041) = 36.91, p < .001, *η^2^_p_* = .569, where post hoc tests showed significant differences across all question combinations (p*_bonferroni_* < .001). These results partially confirm our initial hypothesis. Please refer to the graph in Fig 1 for a representation of the response latencies of the two groups across the three question conditions.

**Fig 1.**
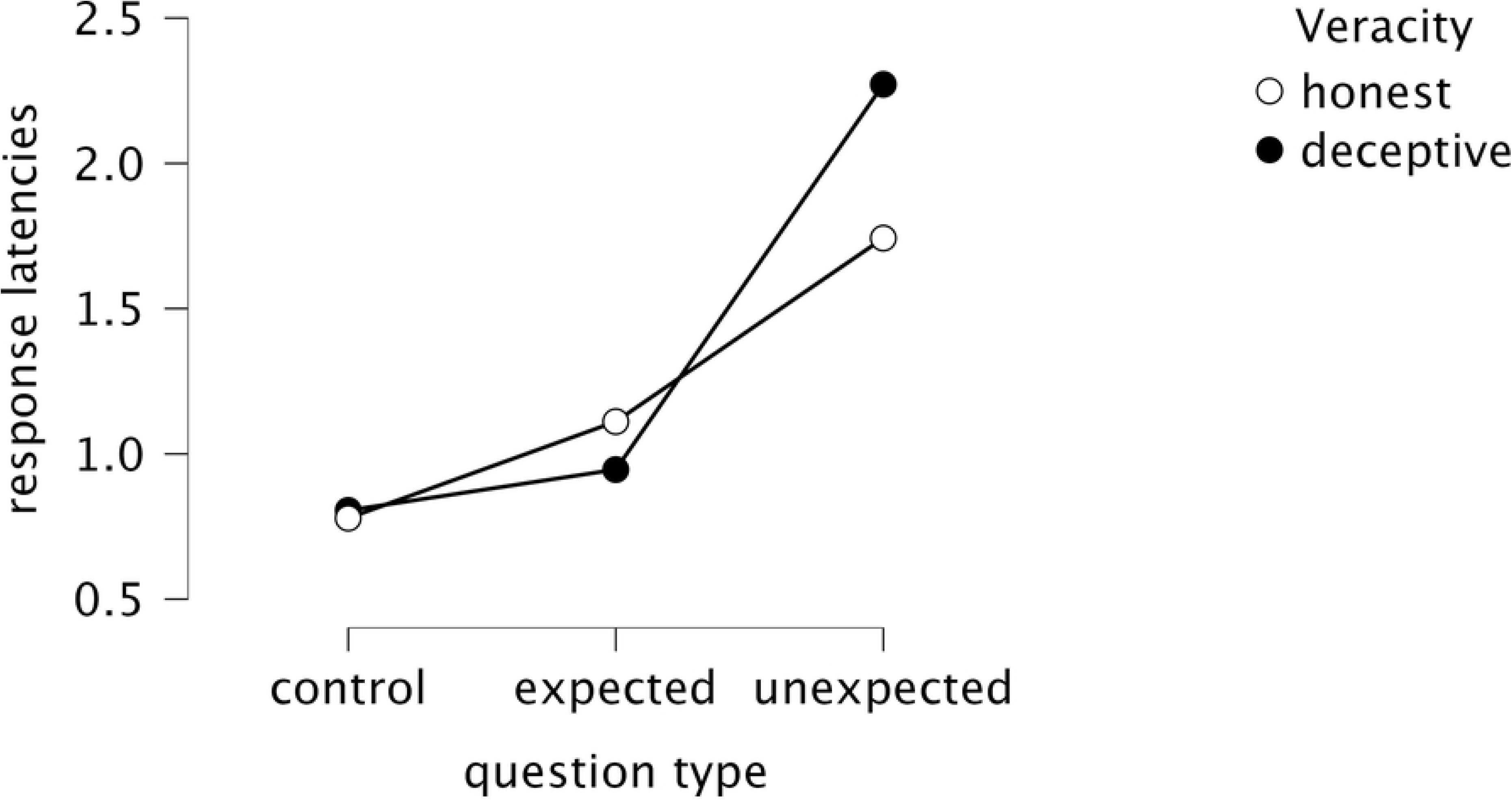
Average response latencies. Average response latencies of the two groups across the three question conditions.

### Analysis of error rate

A 2 (veracity: honest vs. deceptive) x 3 (question type: control vs. expected vs. unexpected) mixed ANOVA was conducted to examine the impact of veracity and question type on the error rate, using the Greenhouse-Geisser correction. The analysis indicated a significant main effect for question type, F*_GG_*(1.463, 84.868) = 69.95, p < .001, *η^2^_p_* = .547, for veracity, F (1,58) = 43.12, p < .001, *η^2^_p_* = 0.426, and for their interaction, F*_GG_*(1.463, 84.868) = 34.38, p < .001, *η^2^_p_* = .372, suggesting that both factors and their interaction are predictors of the error rate. Between-subjects analysis was conducted using an independent samples t-test (Table 2). This revealed a significant difference between the two groups in the error rate associated with unexpected questions, t*_welch_*(42.90) = −7.469, p < .001, d = −1.909, thereby confirming our hypothesis.

**Table 2.** Independent samples t-tests and descriptive statistics of error rates.

|  |  |  |  |  |  | Honest |  | Deceptive |  |
|---|---|---|---|---|---|---|---|---|---|
| Question type | Test | Statistic | df | p | Cohen’s d | M | SD | M | SD |
| Control | Student | -1.601 | 58 | .115 | -0.414 | 0.020 | 0.036 | 0.038 | 0.047 |
|  | Welch* | -1.615 | 55.90 | .112 | -0.415 |  |  |  |  |
| Expected | Student* | -1.046 | 58 | .300 | -0.270 | 0.055 | 0.084 | 0.078 | 0.089 |
|  | Welch | -1.048 | 57.98 | .299 | -0.270 |  |  |  |  |
| Unexpected | Student | -7.307 | 58 | < .001 | -1.888 | 0.089 | 0.086 | 0.336 | 0.186 |
|  | Welch* | -7.469 | 42.90 | < .001 | -1.909 |  |  |  |  |

Furthermore, repeated-measures ANOVAs conducted separately on the groups confirmed significant differences based on question type for both the deceptive, F*_GG_*(1.386, 41.578) = 72.34, p < .001, *η^2^_p_* = .707, and the honest participants, F (1.649, 46.179) = 6.826, p = .004, *η^2^_p_* = .196. Among the deceptive group, post hoc comparisons revealed that the error rate differed significantly between unexpected and control questions (p*_bonferroni_* < .001) and between unexpected and expected questions (p*_bonferroni_* < .001), whilst no difference emerged between control and expected questions (p*_bonferroni_*= .078). In the honest group, a significant difference was found control and unexpected questions (p*_bonferroni_* < .004), no differences emerged between control and expected questions (p*_bonferroni_* = .061) and between expected and unexpected questions (p*_bonferroni_* = .370). The analyses conducted partially confirm our hypothesis. For a representation of the error rates of the two groups across the three question conditions, please refer to the graph in Fig 2.

**Fig 2.**
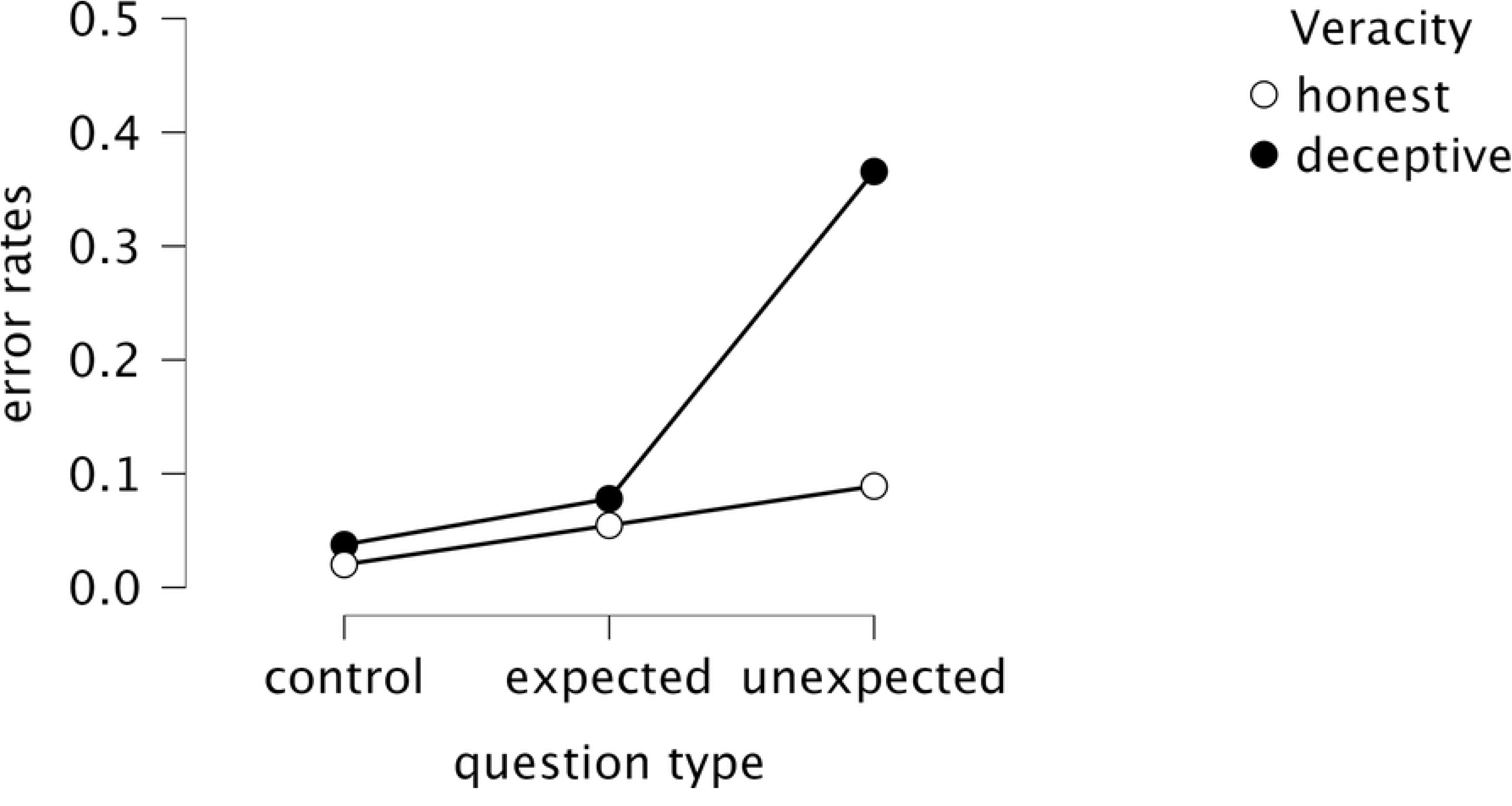
Average error rates. Average error rates of the two groups across the three question types.

### Analysis of inaccuracy rate

To assess the impact of veracity and question type on the inaccuracy rate, a 2 (veracity: honest vs. deceptive) × 3 (question type: control vs. expected vs. unexpected) mixed ANOVA was performed. The results, with Greenhouse-Geisser correction, revealed a significant main effect of question type, F*_GG_*(1.099, 63.744) = 102.19, p < .001, *η^2^_p_* = .638, of veracity, F (1, 58) = 36.25, p < .001, *η^2^_p_* = .385, and a significant interaction between these, F*_GG_*(1.099, 63.744) = 35.70, p < .001, *η^2^_p_* = .381. This suggests that the question type, the veracity and the interaction between these factors are predictors of the inaccuracy rate. The independent samples t-test (Table 3) confirmed a significant difference in the inaccuracy rate for unexpected questions between the two groups, t*_welch_*(48.21) = −6.285, p < .001, d = −1.610. These results confirm the initial hypothesis, showing that the unexpected questions increased inaccuracy rates overall, with a significantly greater effect in the deceptive group.

**Table 3.**
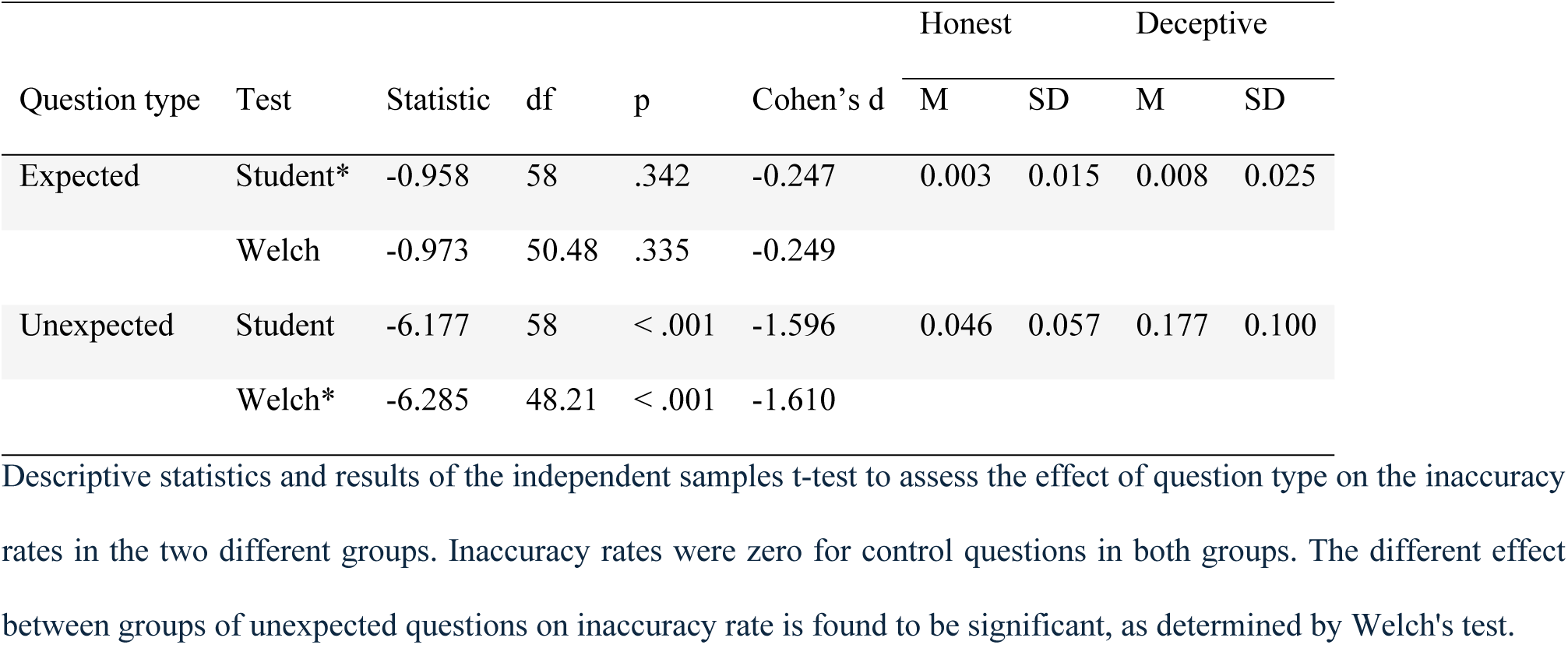
Independent samples t-tests and descriptive statistics of inaccuracy rates.

|  |  |  |  |  |  | Honest |  | Deceptive |  |
|---|---|---|---|---|---|---|---|---|---|
| Question type | Test | Statistic | df | p | Cohen’s d | M | SD | M | SD |
| Expected | Student* | -0.958 | 58 | .342 | -0.247 | 0.003 | 0.015 | 0.008 | 0.025 |
|  | Welch | -0.973 | 50.48 | .335 | -0.249 |  |  |  |  |
| Unexpected | Student | -6.177 | 58 | < .001 | -1.596 | 0.046 | 0.057 | 0.177 | 0.100 |
|  | Welch* | -6.285 | 48.21 | < .001 | -1.610 |  |  |  |  |

In addition, two separate repeated-measures ANOVAs were conducted to examine the effect of question type within each group. The Greenhouse-Geisser correction was applied in the analyses. Both in the deceptive group, F*_GG_*(1.093, 32.780) = 88.47, p < .001, *η^2^_p_* = .747, and in the honest group, F*_GG_*(1.110, 31.074) = 18.34, p < .001, *η^2^_p_* = .396, the results revealed a significant effect of question type. In the deceptive group, a post hoc analysis revealed significant differences in the inaccuracy rate between unexpected and control questions (p*_bonferroni_* < .001) and between unexpected and expected questions (p*_bonferroni_* < .001), whilst no difference emerged between control and expected questions (p*_bonferroni_* = .249). The same pattern was observed in the honest group, where post hoc tests showed significant differences in the unexpected-control questions combinations (p*_bonferroni_* < .001) and unexpected-expected questions combinations (p*_bonferroni_*< .001), whilst no significant difference was found in the control-expected questions combination (p*_bonferroni_* = .978). These results confirm our initial hypothesis. For a graphical representation of the inaccuracy rate of the two groups across the three question conditions, please refer to Fig 3.

**Fig 3.**
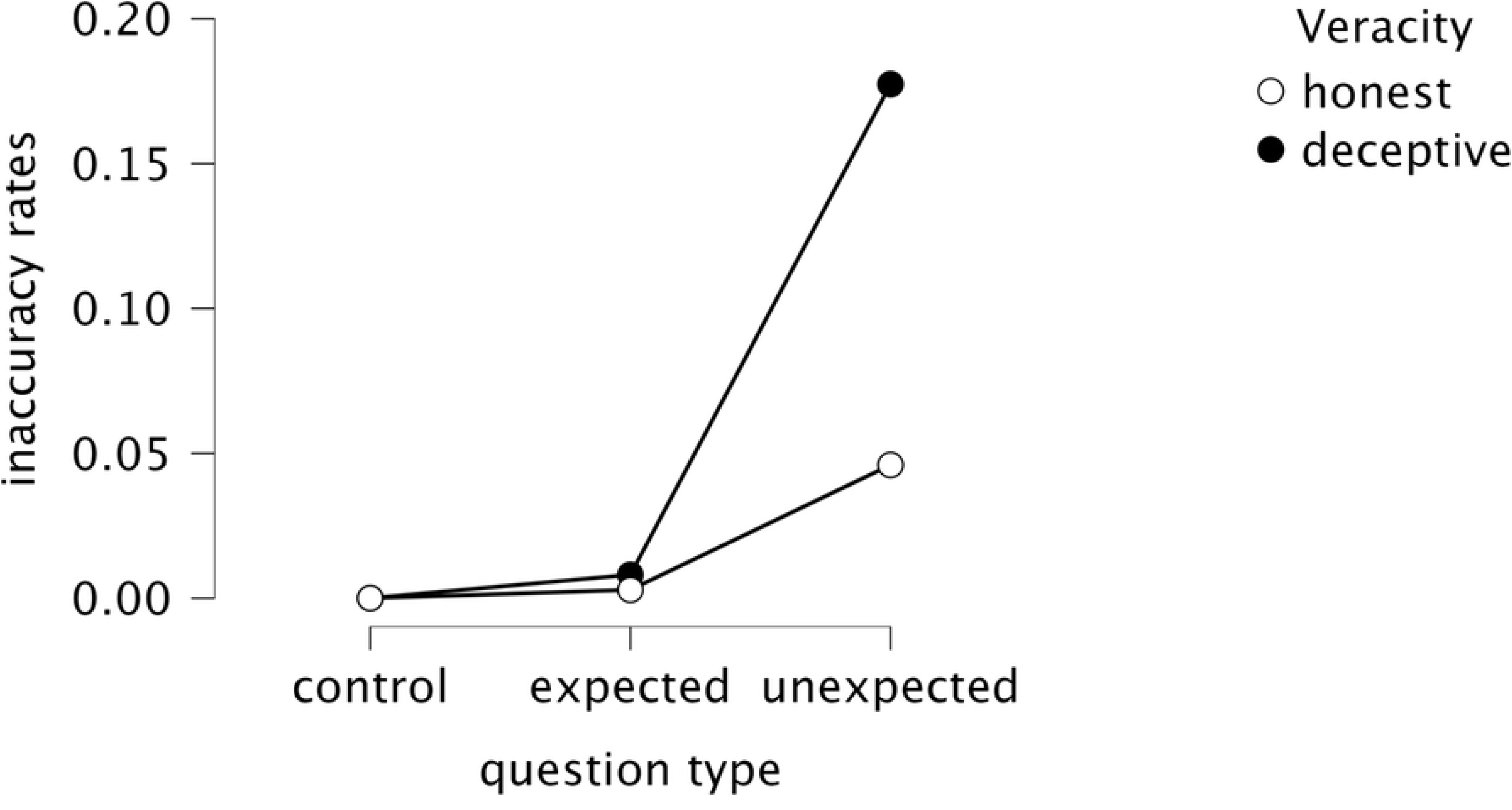
Average inaccuracy rates. Average inaccuracy rates of the two groups across the three question types.

### Analysis of perceived difficulty

For the analysis of perceived difficulty, a 2 (veracity: honest vs deceptive) x 3 (question type: control vs expected vs unexpected) mixed ANOVA with Greenhouse-Geisser correction was conducted. This revealed a significant main effect for question type, F*_GG_*(1.663, 96.461) = 172.97, p < .001, *η^2^_p_* = .749, and a significant interaction between veracity and question type, F*_GG_*(1.663, 96.461) = 18.61, p < .001, *η^2^_p_* = .243. The main effect of veracity was not significant, F (1.663, 96.461) = 3.707, p = .059 *η^2^_p_* = .060. The independent samples t-test indicated that deceptive participants perceive unexpected questions as significantly more difficult than honest participants, t(58) = −3.718, p < .001, d = −0.960. These results confirm our hypothesis, as on average, the deceptive subjects rated answering unexpected questions as more difficult than the honest subjects (Table 4).

**Table 4.**
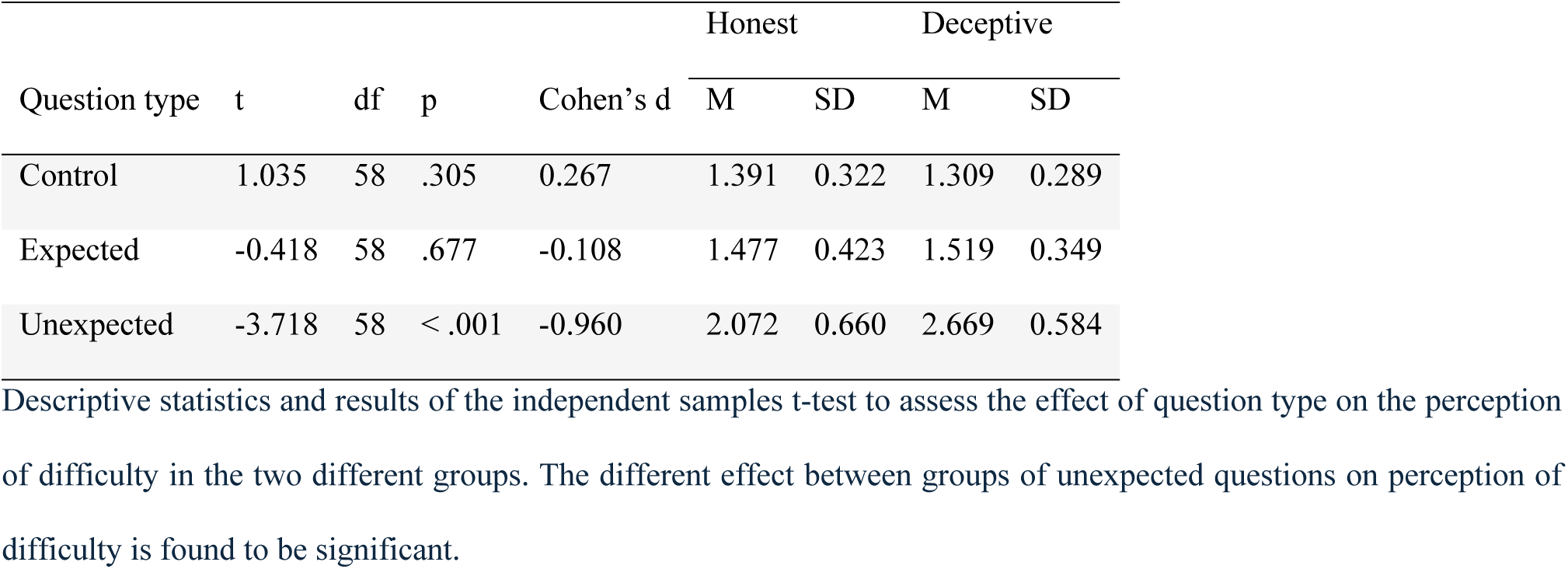
Independent samples t-tests and descriptive statistics of perceived difficulty.

### Analysis of perceived predictability

A 2 (veracity: honest vs deceptive) x 3 (question type: control vs expected vs unexpected) mixed ANOVA, with Greenhouse-Geisser correction, on the perception of predictability revealed a significant main effect of question type, F*_GG_*(1.304, 75.650) = 77.994, p < .001, *η^2^_p_* = .574, whilst no significant effects related to veracity (p = .522) nor an interaction between the two factors (p = .418) emerged. Expected questions were rated as significantly more predictable than unexpected ones for both groups (Table 5), confirming the manipulation of the variable.

**Table 5.** Independent samples t-tests and descriptive statistics of perceived predictability.

|  |  |  |  |  | Honest |  | Deceptive |  |
|---|---|---|---|---|---|---|---|---|
| Question type | t | df | p | Cohen’s d | M | SD | M | SD |
| Control | -1.336 | 58 | .177 | -0.353 | 1.784 | 0.865 | 2.175 | 1.291 |
| Expected | -0.959 | 58 | .342 | -0.248 | 3.718 | 0.752 | 3.917 | 0.843 |
| Unexpected | 1.332 | 58 | .188 | 0.344 | 3.032 | 0.825 | 2.777 | 0.651 |

### Classification of individual subjects

To assess the possibility of correctly classifying an individual participant into one of the two categories (honest or deceptive) based on their responses, two machine-learning models were conducted using the modelling procedure described in the Method section.

Table 6 shows the results of the two algorithms, identifying the random forest as the best-performing algorithm with a discriminative power of 96.7% (AUC = 0.967) (Fig 4), whereas cross-validated accuracy was 0.900 (0.933 at Youden’s cut-off). Further analyses were conducted to identify the error rate at Youden’s cut-off (Table 7): across the out-of-fold predictions of the random forest model, there was a single false negative and 3 false positives, which identifies the model as particularly sensitive in classifying deceptive participants.

**Fig 4.**
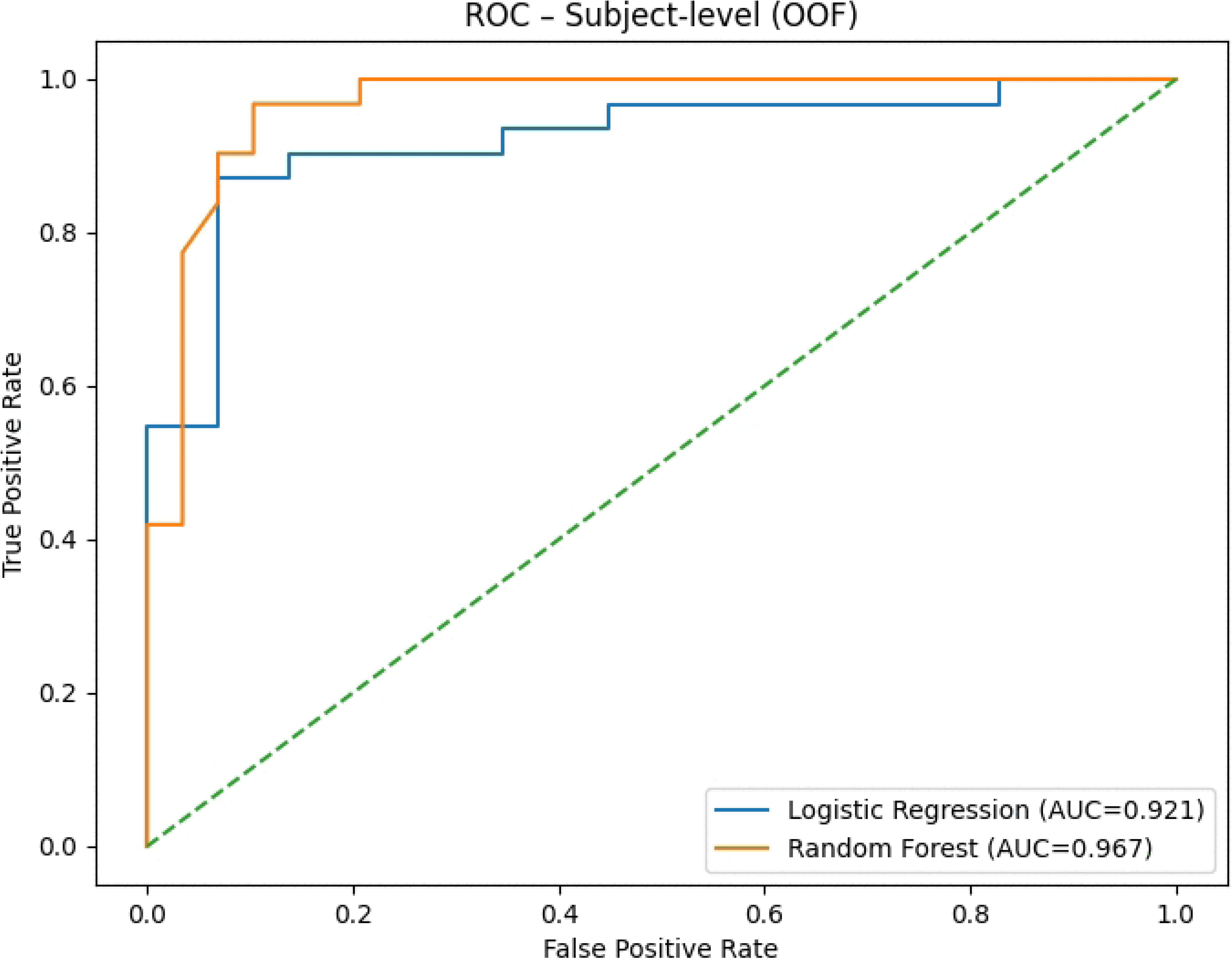
Individual subject classification ROC curves. ROC curves for the logistic regression model (blue) and the random forest model (orange).

**Table 6.** Performance metrics for the two algorithms in classifying honest and deceptive participants.

| **Model** | **Performance** |  |  |  |  |  |  |
|---|---|---|---|---|---|---|---|
|  | Accuracy | Accuracy @ cut-off | AUC | AUC (OOF) | Precision | Recall | F1 |
| Logistic regression | 0.883 | 0.900 | 0.911 | 0.921 | 0.867 | 0.900 | 0.878 |
| Random forest | 0.900 | 0.933 | 0.967 | 0.967 | 0.908 | 0.900 | 0.899 |

The metrics shown in the table are: accuracy, accuracy at Youden cut-off, area under the curve (AUC), area under the curve out-of-fold (AUC (OOF)), precision, recall, and F1. Accuracy, AUC, precision, recall and F1 are 10-fold cross-validation averages at the default probability threshold (0.50), whereas “Accuracy @ cut-off” refers to the confusion matrix at Youden’s cut-off (Table 7); AUC (OOF) is the AUC of the pooled out-of-fold predictions.

**Table 7.**
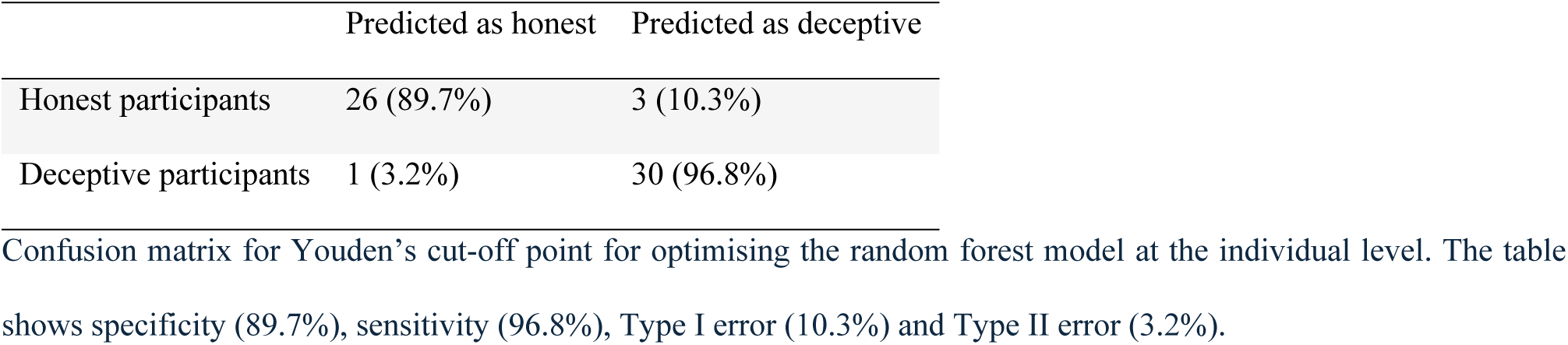
Individual subject classification confusion matrix.

|  | Predicted as honest | Predicted as deceptive |
|---|---|---|
| Honest participants | 26 (89.7%) | 3 (10.3%) |
| Deceptive participants | 1 (3.2%) | 30 (96.8%) |

### Classification of individual items

The analyses classifying individual responses as honest or deceptive were conducted in a similar manner to the previous ones, but using response latency, error and inaccuracy rates of the individual response as classification indicators, rather than considering the entire set of questions. The logistic regression model performed best, with an accuracy of 70% at Youden’s cut-off (Table 8). Despite this, at that cut-off the model has a high percentage of false positives (17.8%) and, above all, false negatives (41.4%) (Table 9).

**Table 8.** Performance metrics for the two algorithms in classifying individual responses as honest or deceptive.

| **Model** | **Performance** |  |  |  |  |  |  |
|---|---|---|---|---|---|---|---|
|  | Accuracy | Accuracy @ cut-off | AUC | AUC (OOF) | Precision | Recall | F1 |
| Logistic regression | 0.696 | 0.700 | 0.738 | 0.733 | 0.778 | 0.572 | 0.657 |
| Random forest | 0.611 | 0.679 | 0.743 | 0.728 | 0.733 | 0.551 | 0.626 |

The metrics shown in the table are: accuracy, accuracy at Youden cut-off, area under the curve (AUC), area under the curve out-of-fold (AUC (OOF)), precision, recall, and F1. Accuracy, AUC, precision, recall and F1 are 10-fold cross-validation averages at the default probability threshold (0.50), whereas “Accuracy @ cut-off” refers to the confusion matrix at Youden’s cut-off (Table 9); AUC (OOF) is the AUC of the pooled out-of-fold predictions.

**Table 9.**
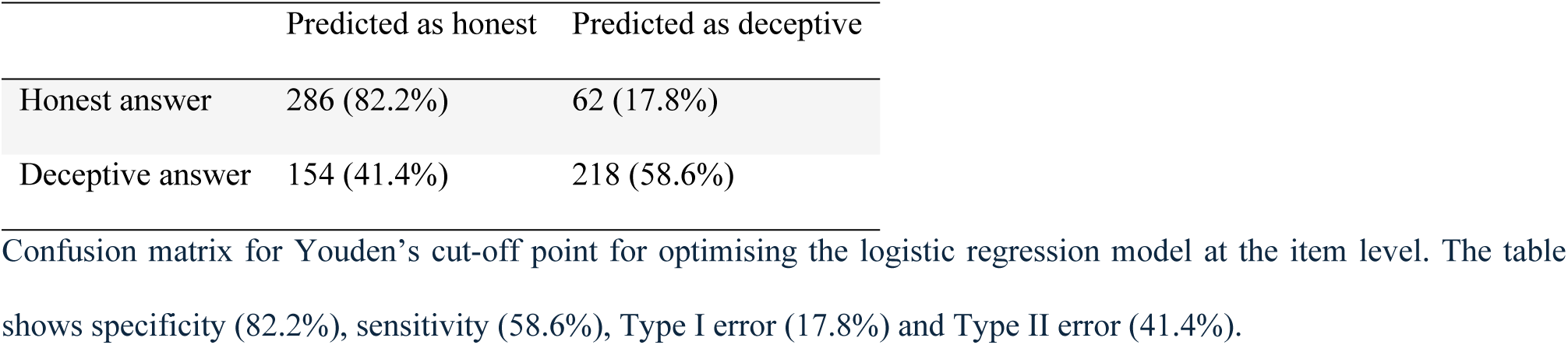
Individual item classification confusion matrix.

|  | Predicted as honest | Predicted as deceptive |
|---|---|---|
| Honest answer | 286 (82.2%) | 62 (17.8%) |
| Deceptive answer | 154 (41.4%) | 218 (58.6%) |

### Cut-off analysis

In order to identify an average threshold for response latency to unexpected questions beyond which the subject can be classified as deceptive, the following formula was applied: Threshold = Mean(Unexpected RT) − [Mean(Control RT) + Mean(Expected RT)] / 2. The latency of each individual unexpected question (RT_Unexpected,i_, *i* = 1, …, 12) was compared with the same subject-specific baseline.

The Youden index (J) was then used to calculate the threshold that optimises sensitivity and specificity. When the average latency to unexpected questions exceeds the subject’s baseline (average latency to control and expected questions) by at least 0.618 seconds, the participant is classified as deceptive. The classification of the subject based on this average-latency indicator yielded an AUC of .742, a sensitivity of 90.3% and a false-positive rate of 44.8%.

When the subject exceeds this threshold in 7 or more of the 12 unexpected questions, classification accuracy rises to 75% but error rates remain high (Table 10).

**Table 10.**
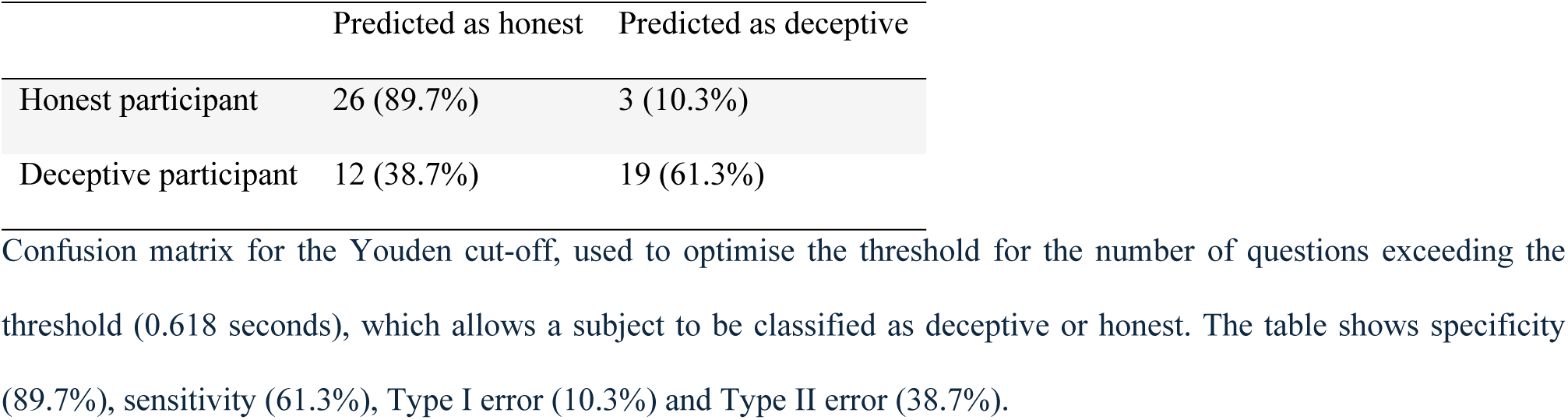
Cut-off confusion matrix for the 7-out-of-12 rule.

|  | Predicted as honest | Predicted as deceptive |
|---|---|---|
| Honest participant | 26 (89.7%) | 3 (10.3%) |
| Deceptive participant | 12 (38.7%) | 19 (61.3%) |

## Discussion

This study investigated response latency, error and inaccuracy rates to unexpected questions as a useful tool to differentiate deceptive from honest participants. Overall, deceptive participants showed larger increases in these indices to unexpected questions than honest participants, extending prior findings in the field.

### Behavioural indicators of cognitive load

The deceptive group showed longer response latencies and higher error and inaccuracy rates to unexpected questions compared to expected and control questions. This pattern corroborates the results of previous studies [18, 19] and is consistent with the hypothesis that the construction of deception, while maintaining narrative coherence, requires greater cognitive load. Similarly, in the honest group, longer response latencies and more errors were observed for unexpected questions compared to expected and control questions, but the difference appears to be significantly smaller than that observed in the deceptive group.

Analyses comparing the two groups revealed that deceptive participants exhibit longer response latencies and higher error and inaccuracy rates to unexpected questions compared to the honest subjects. In the control questions, conversely, the groups did not differ significantly in response latencies, error and inaccuracy rates.

These results are consistent with previous research [18, 19, 33]. Unexpected questions require a greater cognitive load for both groups, as they concern details that require re-processing and retrieval from less active memory traces. The increased indices in deceptive participants are consistent with the view that deception requires additional cognitive processes involved in designing the lie, monitoring plausibility and coherence, and behavioural control. On this basis, the increased response latency is related to the time required to formulate the false response, and the higher error rate is attributable to reduced control over the response due to cognitive overload. Inaccuracy, on the other hand, may constitute a strategy to reduce the cognitive load, allowing participants to avoid details that could generate contradictions or errors.

An alternative explanation for the elevated response latency and error rates is that memorising an unfamiliar story may itself increase cognitive load. However, the fact that the between-group differences were concentrated in the unexpected questions (significant veracity × question type interactions), rather than spread across all question types, speaks against a generic effect of memorisation difficulty, which would also be expected to affect responses to the expected questions. This argument is not conclusive, as a structural asymmetry remains between querying a lived episodic memory and a brief memorised text, which had been rehearsed to an error-free criterion and therefore made the expected questions highly accessible. The task asymmetry therefore remains a limitation (see below).

An opposite trend was observed in response latency to expected questions, with honest participants responding more slowly than deceptive ones, unlike previous findings [18]. Although this difference was not found to be significant, it may reflect a genuine retrieval challenge: while deceptive subjects memorised a story with fixed details, honest participants provided a free-form text without time-stamped information, making specific questions (e.g. *‘What time did you leave to go home?’*) more difficult to answer.

### Perceived difficulty and predictability

The final questionnaires, administered to assess the perception of the questions, are also of interest. No significant differences emerged regarding the perceived predictability of the questions between the two groups: both honest and deceptive participants rated unexpected questions as less predictable than expected questions. On the other hand, both groups perceived the control questions as unpredictable, as they concerned topics unrelated to the day trip, such as the interview setting and physical characteristics. This result is consistent with the research design and the manipulation of the variable; indeed, the unexpected questions were formulated to take the participant by surprise, so that they could not rely on an answer prepared in advance.

Regarding perceived difficulty, however, the groups agreed that answering unexpected questions required greater effort than answering expected or control questions. Specifically, the deceptive group rated these questions as more complex than honest participants. This result, consistent with previous research [18], likely reflects the different information accessibility: in the honest group, the requested information was less readily accessible. At the same time, the higher difficulty ratings reported by deceptive participants are in line with the additional demands of generating and monitoring a false response.

### Machine-learning and cut-off analysis

Traditional statistical methods focus on differences between groups, failing to provide the necessary precision in the analysis of individual cases, which is essential within the field of application of this study. Therefore, machine-learning models may help extend the analysis toward individual-level prediction, which is relevant to potential forensic and investigative applications. This methodology is widely used in the relevant literature [18, 19]. An innovative aspect of the study was the exploratory identification of cut-off points that may help to take further steps toward identifying a replicable, operational decision-making rule applicable in investigative contexts.

At the individual level, the machine-learning results showed a promising ability to identify a single subject as honest or deceptive within the present sample, based on their response latency, error rate, and inaccuracy rate. These results require replication and external validation before individual-level application can be considered. Both logistic regression and random forest achieved high classification accuracy (up to 93.3%) and good discriminative power (AUC = .967). On the other hand, the models constructed at the item level were designed to assess the possibility of classifying a subject as deceptive or honest based on a single response to an unexpected question. This analysis was intended to explore whether the single response provided by the subject to an unexpected question could be classified as true or false. Although these machine-learning algorithms produced moderate results, with an accuracy of 70% and an AUC of 0.738, they are subject to a high error rate, particularly false negatives (41.4%). Consequently, these models are not sufficiently reliable for forensic application or for overcoming the threshold of *“reasonable doubt”*.

The threshold analysis of response latency to unexpected questions revealed that when the average latency to unexpected questions exceeded the subject’s baseline by at least 0.618 seconds, the model classified the participant as deceptive. The threshold identified showed moderate discriminatory power (AUC = .742) and high sensitivity (90.3%), but was associated with a high percentage of false positives (44.8%), limiting its practical utility. Furthermore, a threshold number of responses to unexpected questions was determined that must exceed the cut-off by at least 0.618 seconds compared to the baseline to classify the subject as deceptive. The threshold identified, of 7 out of 12 responses, led to an increase in accuracy (75%), a moderate result but characterised by a high percentage of false negatives (38.7%).

A comparison between the subject level and the item level reveals a marked loss of discriminatory power when the analysis is conducted on a single response, rather than considering the overall pattern of responses. Similar results were found in the study conducted by Melis and colleagues in 2024 [18], suggesting that the use of a greater amount of information reduces intra-individual variability. The application of the response latency threshold to multiple responses further corroborates this increased classification accuracy when considering the whole response pattern. This aligns with the cognitive load theory in which the single answer may not reveal greater cognitive load, but the entire set of responses highlights detectable patterns. The results are promising, but their practical application remains premature at the present stage.

### Limitations and future prospects

Although the findings provide relevant information regarding the role of response latency, error and inaccuracy rates to unexpected questions as a method for detecting deception, this study has certain limitations that are worth discussing.

Firstly, it is important to highlight the absence of both a double-blind procedure and an inter-rater reliability assessment: the entire process of conducting the interviews and analysing the audio recordings was entrusted to a single experimenter. This decision, dictated by organisational constraints, may have introduced confirmation bias in the categorisation of responses. The adoption of a double-blind experimental design, in which the interviewer and the coder are independent and unaware of participants’ veracity condition, would minimise expectancy effects. Furthermore, future studies should have a second rater, blinded to veracity and to the hypotheses, re-code a random subset of the recordings and report inter-rater reliability.

An additional limitation is the reduced control over the accuracy of the responses provided by the honest participants, due to the use of free-form narration of the event under analysis. This choice, which allowed greater ecological validity of the research, made it difficult to clearly distinguish memory errors, inaccuracies, and genuinely accurate responses, potentially inflating variability in the honesty condition. Moreover, neither the retention interval since the honest participants’ trip nor the length and level of detail of their free-form accounts were standardised, whereas the memorised story had a fixed length and level of detail. Furthermore, another limitation of the experimental design worth addressing concerns the construction of the tasks for the two experimental groups: recalling the most recent day trip for the honest group and memorising a narrative in just five minutes for the deceptive group. Although this is an experimental design commonly found in the literature [18], it entails cognitively different tasks for the two groups. Given the highly specific information requested, the results may partly reflect the different familiarity with the information, not only the cognitive demands of deception. To verify the actual accuracy of this study, future research could introduce an additional control condition, in which the honest participants are given a recent memorisation task before the interview, for example, the story of an activity performed shortly beforehand.

Furthermore, the composition of the sample, consisting mainly of university students, limits the generalisability of the results to the general population. Future studies should therefore be based on more heterogeneous samples to verify the robustness of the results across different populations.

This study represents a low-stakes context in which the motivation to lie is lower, as the consequences are less severe than in real investigative contexts. Under these conditions, participants may therefore have employed less elaborate and less convincing lying strategies than those used in real high-stakes situations. The imposition of cognitive load through task-switching is a valid tool for investigating deception in low-stakes contexts, allowing high-stakes contexts to be simulated and thereby increasing the ecological validity of the study. Nevertheless, further research in real-world contexts is required.

A key strength of the research, in addition to the study of deception in the recounting of autobiographical events, was the expansion of response categories through the introduction of inaccuracy. This allowed, at least in part, for the overcoming of the dichotomy between “bald-faced lies” (100% false statement) and “bald-faced truths” (100% true statement), criticised by McCornack and colleagues [35, 36]. However, future research could investigate scenarios in which deceptive participants alter only certain details of the event being recounted, as frequently occurs, rather than constructing entirely fictional narratives.

## Conclusions

The results of this study support the value of response latency, error rate, and inaccuracy rate to unexpected questions as promising indicators of deception in autobiographical event narratives. Crucially, these findings reflect deception as operationalized under a specific experimental paradigm (comparing episodic retrieval to recent memorization), rather than examining deception *per se*. At the same time, the use of machine-learning models and the identification of specific cut-off points may facilitate the transformation of these implicit indicators into practical decision-making criteria for investigative contexts.

On the practical side, our study highlighted the potential usefulness of the unexpected question technique for investigative work as response latencies can be extracted post hoc from audio recordings without alerting the interviewee to the diagnostic purpose. This applicability should not be overstated, however. Response latency may therefore offer a particularly practical indicator; by contrast, classifying answers as erroneous or inaccurate requires a verified reference account against which they can be assessed, which is rarely accessible in real investigations. Traditional lie detection techniques are awareness-based methods, meaning the subject under examination is fully aware that their credibility is being assessed. This distinction is particularly relevant in investigative settings, where the ability to conceal the diagnostic purpose of an interview may be advantageous. On the other hand, the unexpected questions technique is a relatively non-invasive and implicit strategy, able to obscure the diagnostic purpose of the interview, thereby reducing opportunities for strategic self-monitoring by the respondent.

In addition to overcoming the primary limitations of traditional lie detection techniques, this study partially addresses the ecological validity gap identified in previous research. By applying the unexpected questions technique to event narratives and introducing inaccuracy rate, this research points toward more operationally realistic lie detection paradigms, while still requiring replications under more field-based and methodologically controlled conditions.

## Acknowledgments

An AI-assisted critical review was conducted on an earlier version of the manuscript using Google Gemini’s Deep Research. The tool was used to identify potential issues in the manuscript‘s clarity and argumentation and to suggest improvements. The AI tool was not used for data collection, statistical analyses, or the generation of research results. The suggestions were critically evaluated by the authors, who retained full responsibility for the revisions and the final content of the manuscript.

## Supporting Information

**S1 Table 4. Independent sample t-tests and descriptive statistics of response latencies.** Descriptive statistics and results of the independent samples t-test to assess the effect of question type on response latencies in the two different groups. The different effect between groups of unexpected questions on response latency is found to be significant, as determined by Student’s t-test.

**S2 Table 2. Independent sample t-tests and descriptive statistics of error rates.** Descriptive statistics and results of the independent samples t-test to assess the effect of question type on the error rates in the two different groups. The different effect between groups of unexpected questions on error rate is found to be significant, as determined by Welch’s test.

**S3 Table 3. Independent sample t-tests and descriptive statistics of inaccuracy rates.** Descriptive statistics and results of the independent samples t-test to assess the effect of question type on the inaccuracy rates in the two different groups. Inaccuracy rates were zero for control questions in both groups. The different effect between groups of unexpected questions on inaccuracy rate is found to be significant, as determined by Welch’s test.

**S4 Table 4. Independent sample t-tests and descriptive statistics of perceived difficulty.** Descriptive statistics and results of the independent samples t-test to assess the effect of question type on the perception of difficulty in the two different groups. The different effect between groups of unexpected questions on perception of difficulty is found to be significant.

**S5 Table 5. Independent sample t-tests and descriptive statistics of perceived predictability.** Descriptive statistics and results of the independent samples t-test to assess the effect of question type on the perception of predictability in the two different groups. No significant results were found.

**S6 Table 5. Performance metrics for the two algorithms in classifying honest and deceptive participants.** The metrics shown in the table are: accuracy, accuracy at Youden cut-off, area under the curve (AUC), area under the curve out-of-fold (AUC (OOF)), precision, recall, and F1. Accuracy, AUC, precision, recall and F1 are 10-fold cross-validation averages at the default probability threshold (0.50), whereas “Accuracy @ cut-off” refers to the confusion matrix at Youden’s cut-off (Table 7); AUC (OOF) is the AUC of the pooled out-of-fold predictions.

**S7 Table 7. Individual subject classification confusion matrix**. Confusion matrix for Youden’s cut-off point for optimising the random forest model at the individual level. The table shows specificity (89.7%), sensitivity (96.8%), Type I error (10.3%) and Type II error (3.2%).

**S8 Table 8. Performance metrics for the two algorithms in classifying individual responses as honest or deceptive.** The metrics shown in the table are: accuracy, accuracy at Youden cut-off, area under the curve (AUC), area under the curve out-of-fold (AUC (OOF)), precision, recall, and F1. Accuracy, AUC, precision, recall and F1 are 10-fold cross-validation averages at the default probability threshold (0.50), whereas “Accuracy @ cut-off” refers to the confusion matrix at Youden’s cut-off (Table 9); AUC (OOF) is the AUC of the pooled out-of-fold predictions.

**S9 Table 9. Individual item classification confusion matrix.** Confusion matrix for Youden’s cut-off point for optimising the logistic regression model at the item level. The table shows specificity (82.2%), sensitivity (58.6%), Type I error (17.8%) and Type II error (41.4%).

**S10 Table 6. Cut-off confusion matrix for the 7-out-of-12 rule.** Confusion matrix for the Youden cut-off, used to optimise the threshold for the number of questions exceeding the threshold (0.618 seconds), which allows a subject to be classified as deceptive or honest. The table shows specificity (89.7%), sensitivity (61.3%), Type I error (10.3%) and Type II error (38.7%).

**S11 Fig 5. Average response latencies.** Average response latencies of the two groups across the three question conditions.

**S12 Fig 6. Average error rates**. Average error rates of the two groups across the three question types.

**S13 Fig 7. Average inaccuracy rates.** Average inaccuracy rates of the two groups across the three question types.

**S14 Fig 8. Individual subject classification ROC curves.** ROC curves for the logistic regression model (blue) and the random forest model (orange).

**S15 File 1. Supplementary materials.**

